# Balancing Prior Knowledge and Sensory Data in a Predictive Coding Model: Insights into Coherent Motion Detection in Schizophrenia

**DOI:** 10.1101/2024.05.28.596140

**Authors:** Elnaz Nemati, David B. Grayden, Anthony N. Burkitt, Parvin Zarei Eskikand

## Abstract

This study introduces a biologically plausible computational model based on the predictive coding algorithm, providing insights into motion detection processes and potential deficiencies in schizophrenia. The model decomposes motion structures into individual and shared sources, highlighting a critical role of surround suppression in detecting global motion. This biologically plausible model sheds light on how the brain extracts the structure of motion and comprehends shared or coherent motion within the visual field. The results obtained from random dot stimuli underscore the delicate balance between sensory data and prior knowledge in coherent motion detection. Model testing across varying noise levels reveals longer convergence times with higher noise, consistent with psychophysical experiments showing that response duration (e.g., reaction time or decision-making time) also increases with noise levels. The model suggests that an excessive emphasis on prior knowledge extends the convergence time in motion detection. Conversely, for faster convergence, the model requires a certain level of prior knowledge to prevent excessive disturbance due to noise. These findings contribute to potential explanations for motion detection deficiencies observed in schizophrenia.

## 1 Introduction

In a world where visual information is constantly changing and can be ambiguous, efficiently understanding our environment necessitates the brain to discern what motion occurs from the visual input received in the environment, including both self-motion and the motion of other objects. In understanding the complex interplay between sensory input and neural processing during motion perception, animal models provide valuable insights into the underlying neural circuitry. For instance, the comprehensive review by Gollisch and Meister (2010) highlights the computational capabilities of neural circuits in the early visual areas, revealing mechanisms by which these circuits process motion efficiently.

Motion detection is a complex perceptual process that involves integrating prior knowledge, namely past visual experiences, with sensory data, namely the current visual information. This process is not performed in isolation. Our visual system considers our self-motion and the relative motion and interaction of neighboring objects (Born, 2000; Braddick, 1993), which play crucial roles in how motion stimuli are identified and interpreted. The activity of the neurons in the visual pathway is strongly influenced by the motion in the surround of their receptive fields (Allman et al., 1985; Bradley and Andersen, 1998; Tadin et al., 2003). Nearby stimuli shape the detection and comprehension of coherent motion. (Braddick, 1993; Born, 2000; Tadin et al., 2003). The complex microcircuits of the brain that are learned through this process allow the seamless integration of past and present motion information, enabling effective navigation in the environment.

### 1.1 Exploring Hierarchical Motion Perception: From Decomposition Principles to Predictive Coding Models

The brain achieves motion perception by using statistical relationships in the velocities of objects (Bill et al., 2022) and by breaking down object motions into underlying motion sources. These motion sources may not have a one-to-one relationship with objects. For example, for a school of fish, the movement of the fish can be broken down into both the shared motion of the school and the individual motion of each fish, as illustrated in Figure. 1. Although it may seem abstract in concept, shared motion significantly enhances our perception and understanding of such a visual scene involving many components that display coherent motion. By identifying the underlying motion structure, interpretation of the scene is enhanced, facilitating navigation and prediction tasks.

**Figure 1:**
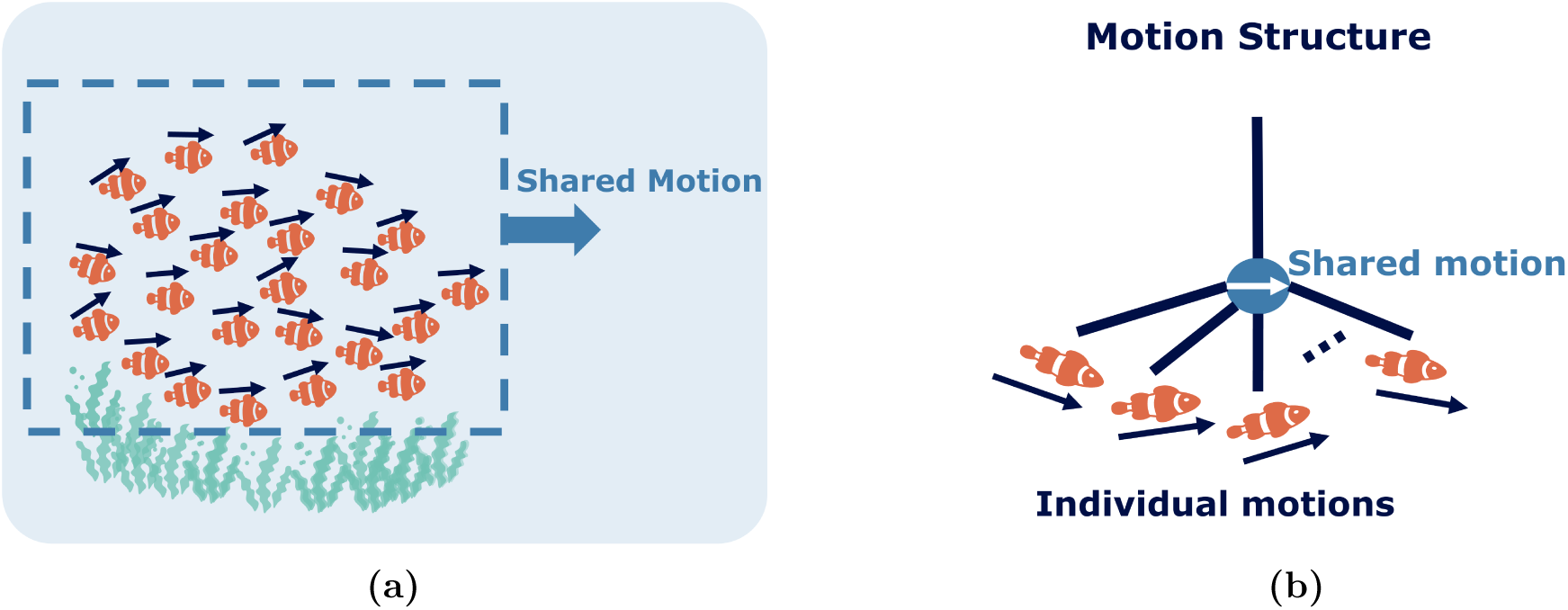
Illustration of visual motion structure separation in complex scenes. (a) A scene of a school of fish, all moving collectively to the right (indicated by the pale blue arrow), with each fish exhibiting its unique motion (indicated by a navy blue arrow above each fish). (b) The scene’s motion can be decomposed into shared motion (averaged over all fish) and individual motion components, relative to the shared motion.

Pioneering studies by Johansson (1950)significantly advanced the exploration of hierarchical motion perception by demonstrating the decomposition of motion through displays of moving dots. These studies uncovered that the simple motion of individual components in a visual scene can evoke complex perceptual experiences. This work supports the concept that the visual system uses vector analysis within a scene, distinguishing between the shared motion of objects and their individual motions. In subsequent studies on biological motion, Johansson (1973) demonstrated that subtracting the shared motion of components from the visual input enables observers to perceive the relative motions of individual parts. This iterative subtraction method enabled a deeper understanding of the multilevel motion structure in visual scenes, laying the groundwork for our current understanding of hierarchical motion perception.

The vector analysis theory in hierarchical motion perception breaks down object motion into common and relative components, offering insightful yet computationally limited perspectives. While it assumes the presence of known motion components and structures, the theory does not inherently detect these elements from sensory data, thus limiting its application in real-world motion analysis (Shum and Wolford, 1983). Complex scenes with multiple possible vector analyses amplify this issue. Researchers have developed several theories to clarify ambiguities in motion interpretation. The ‘minimum principle’ (Restle, 1979) favors the most straightforward interpretations, aligning with Bayesian methods, applicable mainly in ideal, noise-free conditions with specific parametric motions. In contrast, the ‘adjacency principle’ (Gogel, 1974) centers on analyzing cues from relative motion between closely located points. Additionally, the ‘belongingness principle’ (DiVita and Rock, 1997) highlights the effects of coplanarity and reference frames on objects in motion perception.

Recently, researchers using normative models based on Bayesian inference over tree structures have gained insights into more intricate aspects of human motion structure perception. Gershman et al. (2016) introduced a Bayesian framework that uses probabilistic inference over hierarchical motion structures, known as ‘motion trees’, to understand how humans perceive motion structures. Subsequently, Yang et al. (2021) proposed that normative Bayesian models over tree structures could explain crucial aspects of motion perception. These studies primarily focused on modeling motion integration in relation to perceptual outcomes and did not address the complex challenge of real-time motion parsing while concurrently inferring scene structure. Bill et al. (2022) proposed an innovative approach by formulating visual motion perception as online hierarchical inference within a generative model. This continuous-time model effectively explains various aspects of human motion perception and sheds light on the complex interplay between motion parsing and structure inference in real-time perception.

Motion detection heavily relies on past motion information. Traditionally, researchers have overlooked the role of past information cues and prior experience in the study of motion per-ception, presenting a significant gap in the understanding of motion processing. Hierarchical predictive coding offers a promising approach to bridge this gap. This framework posits that the brain constructs a multi-level model, where high-level information influences the processing of sensory data at lower levels (Rao and Ballard, 1999; Millidge et al., 2021). In motion perception, the brain integrates current visual stimuli with past experiences to predict and interpret motion patterns. The brain efficiently and rapidly processes motion by employing predictive coding at lower levels, such as the primary visual cortex (V1) and V2. This method involves the brain comparing top-down predictions from higher-order areas with bottom-up sensory inputs from these lower-order areas, thereby enhancing the understanding of complex motion structures. Predictive coding actively minimizes errors between expected and actual sensory inputs through feedback mechanisms, refining motion perception and integrating it with more complex functions in higher visual areas like the middle temporal (MT) and inferotemporal (IT) cortices. This active processing outlines the neural structures involved and the underlying processes. Despite the theoretical appeal and potential applicability of predictive coding in the domain of motion perception, the literature still shows a scarcity of such models, indicating a valuable direction for future research.

This study aims to explore the brain’s adaptive mechanisms in motion perception through neural connectivity and networks. The central hypothesis of this research is grounded in the concept of ‘predictive coding’ (Rao and Ballard, 1999), where the activity in the higher levels of the hierarchy—such as the medial temporal (MT) area, which represents a higher order within the visual hierarchy relative to the primary visual cortex (V1)—determines (or ‘predicts’) the activity in the lower levels. Although classical predictive coding functions predictively in this hierarchical fashion, it has also been postulated to be utilized by the brain in the temporal domain to forecast future events (Millidge et al., 2024). We test the classical predictive coding model with different stimuli and demonstrate that the model successfully extracts the global motion of the stimuli and replicates the perception observations recorded in psychophysical experiments (Johansson, 1973; Duncker, 1929).

### 1.2 Motion detection deficiency in schizophrenia

Schizophrenia impacts around 24 million individuals globally and is recognized as one of the most debilitating and economically devastating disorders, as reported by the World Health Organization (Institute of Health Metrics and Evaluation, 2021). Individuals diagnosed with schizophrenia have been found to exhibit impairments in tasks related to motion processing, specifically in tasks such as velocity discrimination, where patients seek to identify which stimulus is moving at a faster pace (Chen et al., 1999; Kim et al., 2013). In an fMRI study, it was observed that the Middle Temporal (MT) area in the visual cortex, which is known to be highly specialized for motion detection, is significantly less active in patients with schizophrenia compared to control subjects when conducting motion detection tasks for direction discrimination and speed discrimination (Chen et al., 2008).

A frequently used type of stimulus to investigate the neural circuitry of motion detection is random dots. In the context of schizophrenia, Random Dot Kinematograms (RDKs) prove to be very informative, as they capture the ability of the nervous system to integrate the local motion of individual dots to estimate global motion under different levels of noise. A psychophysical study showed that individuals with schizophrenia require a higher level of coherence in random dots (i.e., less noise) to estimate the global motion of the dots compared to healthy controls (Kim et al., 2013). Furthermore, researchers have observed that patients with schizophrenia show weaker surround suppression in neuronal circuitry for motion detection than their healthy counterparts (Tibber et al., 2013).

This study presents a model to explore the underlying neuronal circuitry associated with the observed motion detection deficiency in schizophrenia. This model synthesizes various factors, notably highlighting the impact of weaker surround suppression in subjects with a motion detection deficiency. We aim to highlight that specific alterations in motion perception, particularly those influenced by surround suppression and prior knowledge weighting, may provide insights into the motion detection impairments commonly observed in schizophrenia. These mechanisms form the basis for explaining why individuals with schizophrenia exhibit prolonged latencies in detecting coherent motion under certain conditions, as demonstrated in psychophysical experiments.

The biological plausibility of our model rests on two key features: (a) it incorporates predictive coding mechanisms that minimize prediction errors through forward and feedback signalling in cortical layers and (b) it includes surround suppression, a well-documented process in the visual cortex that enables the extraction of global motion from complex stimuli. These mechanisms mirror established neural processes in the visual system and allow our model to replicate psychophysical observations, such as longer response times with increased noise levels.

This study is the first to investigate the necessity of balancing sensory input and prior knowledge in detecting the global motion of stimuli and how this balance relates to the observed motion detection deficiency in schizophrenia. Furthermore, despite recent advancements in predictive coding models, no existing model specifically addresses motion detection deficiencies in schizophrenia within the framework of predictive coding. While previous studies have connected predictive coding impairments to broader aspects of schizophrenia such as cognitive and perceptual disruptions (Liddle and Liddle, 2022), our study uniquely focuses on how these impairments manifest specifically in motion detection.

## 2 Materials and Methods

### 2.1 Model Framework Overview

To formalize the process of estimating motion sources for an object, we consider motion dynamics where a set of *M* hidden motion sources influence the velocity of an object, denoted as *v_i_*.

We represent each of these sources as *r_j_* for the *j*^th^ source. The weight *u_i,j_* quantifies the extent to which each motion source affects the velocity of the object *i*. Specifically, we can express the velocity of an object as the sum of the products of the weights and the corresponding motion source contributions.

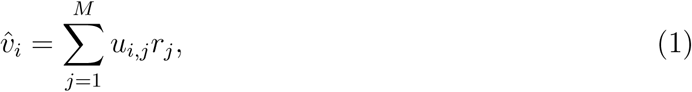

Where *v̂_i_* represents the estimated velocity of the object *i*. The weight *u_i,j_* indicates the influence of the *j*-th motion source on object *i*. These weights are organized into the matrix **U**, and the motion sources *r_j_* are components of the vector **r**, which contains the values for all motion sources. The weights *u_i,j_* range from 0 to 1, where 0 indicates no impact and higher values indicate a stronger influence on the object’s motion. It is important to note that the value of each motion source *r_j_* remains consistent across all objects; only the weight *u_i,j_* varies to reflect the total velocity contribution from each motion source to the object. This framework provides a structured approach for dissecting and analyzing the complex interactions between objects and their motion sources, thereby facilitating a deeper understanding of motion dynamics.

We present a methodology to explain the complexities of motion sources and their structural underpinnings. This approach builds on the classical predictive coding framework (Rao and Ballard, 1999) and incorporates a surround suppression mechanism. This enhancement improves the representation of the interplay between historical motion data and the influence of neighboring objects. The predictive coding framework estimates motion by analyzing neuronal activity within receptive fields and combining it with synaptic weights. Neuronal activity identifies the motion’s source, while synaptic weights quantify how strongly a motion source influences an object’s overall motion. Neurons within receptive fields determine the perceived motion direction, while synaptic weights capture the contribution of each object’s motion to the observed scene dynamics.

We extend the predictive coding model by incorporating surround suppression, a neurobiological mechanism that adjusts neural excitability based on motion context from adjacent stimuli (Tsui et al., 2010). Surround suppression fine-tunes the initial motion predictions that predictive coding generates. Figure. 2 uses a triangle symbol to represent this refinement process. Predictive coding calculates initial estimates as the product of neuronal activities and synaptic weights, **Ur**. Surround suppression evaluates the consistency of motion signals and gates these estimates based on the alignment between center and surround motion.

**Figure 2:**
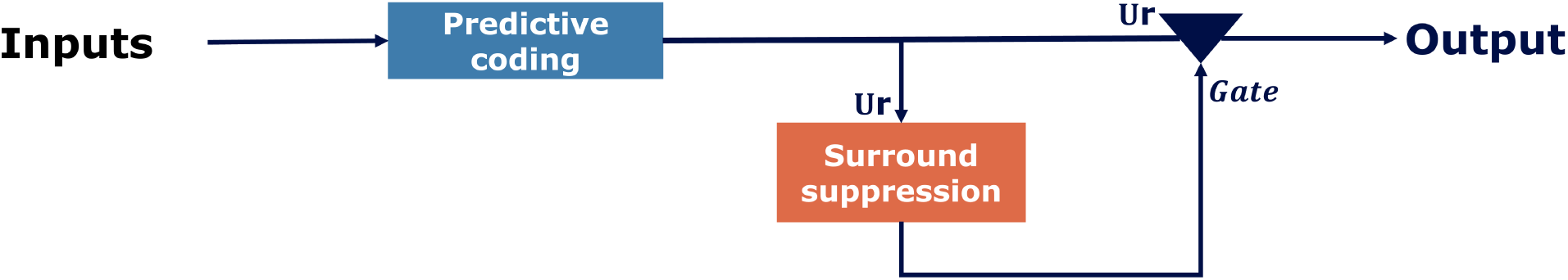
The proposed Motion Estimation Method. The estimated motions generated by the predictive coding, which is the activity of the neurons r multiplied by the synaptic weights **U**, marked on the arrows, are processed by the surround suppression mechanism. Depending on the activation of surround suppression, the estimated motions by the predictive coding are gated and symbolised by a triangle.

Figure. 2 illustrates how the surround suppression mechanism interacts with predictive coding. Surround suppression evaluates motion consistency between the center and surrounding areas and adjusts predictions accordingly. When center and surround motions diverge, surround suppression reduces neural activity to improve the prediction. Predictive coding computes motion estimates across hierarchical levels. It calculates velocity as a weighted sum of item positions, forming the core prediction. Surround suppression compares these velocity predictions (*v* = *Ur*) with the surrounding motion context and uses temporal filters and Heaviside functions to adjust predictions based on the consistency of motion signals.

When initial motion estimates fail to represent an object’s dynamics, the model uses additional predictive coding iterations. This recursive process captures the object’s motion more accurately, refining predictions until they fully represent the observed dynamics.

We systematically examine two-dimensional spatial configurations of objects in visual stimuli using a predictive coding model enhanced with a surround suppression module. This model identifies and quantifies motion sources while considering the influence of adjacent objects. Surround suppression activates when an object’s motion deviates significantly from its surroundings and reduces the impact of these outliers on the motion estimation process. When center and surround motions align, surround suppression remains inactive, allowing accurate motion predictions. This process aligns with findings from Eskikand et al. (2016), showing that suppressive mechanisms enhance motion estimation by adapting neural responses to contextual motion cues.

Incorporating this approach, we are able to extract both the global shared motion and individual motion sources, thereby providing a detailed dissection of motion perception. This methodology, which marries the classical framework of predictive coding (Rao and Ballard, 1999) with a surround suppression mechanism, not only enhances the model’s ability to capture the nuanced interplay between historical motion data and the effects of proximal stimuli but also enables the differentiation between shared and unique motion dynamics, as elaborated in subsequent sections.

### 2.2 Classical Predictive Coding

The predictive coding model offers a compelling framework for understanding brain function, likening it to an algorithm that predicts future events based on sensory data. The brain continuously makes educated guesses about incoming sensory information and adjusts these predictions as new data arrives. This model organizes its processes in a multilevel structure, as shown in Figure. 3a, with each level playing a unique role in processing information.

**Figure 3:**
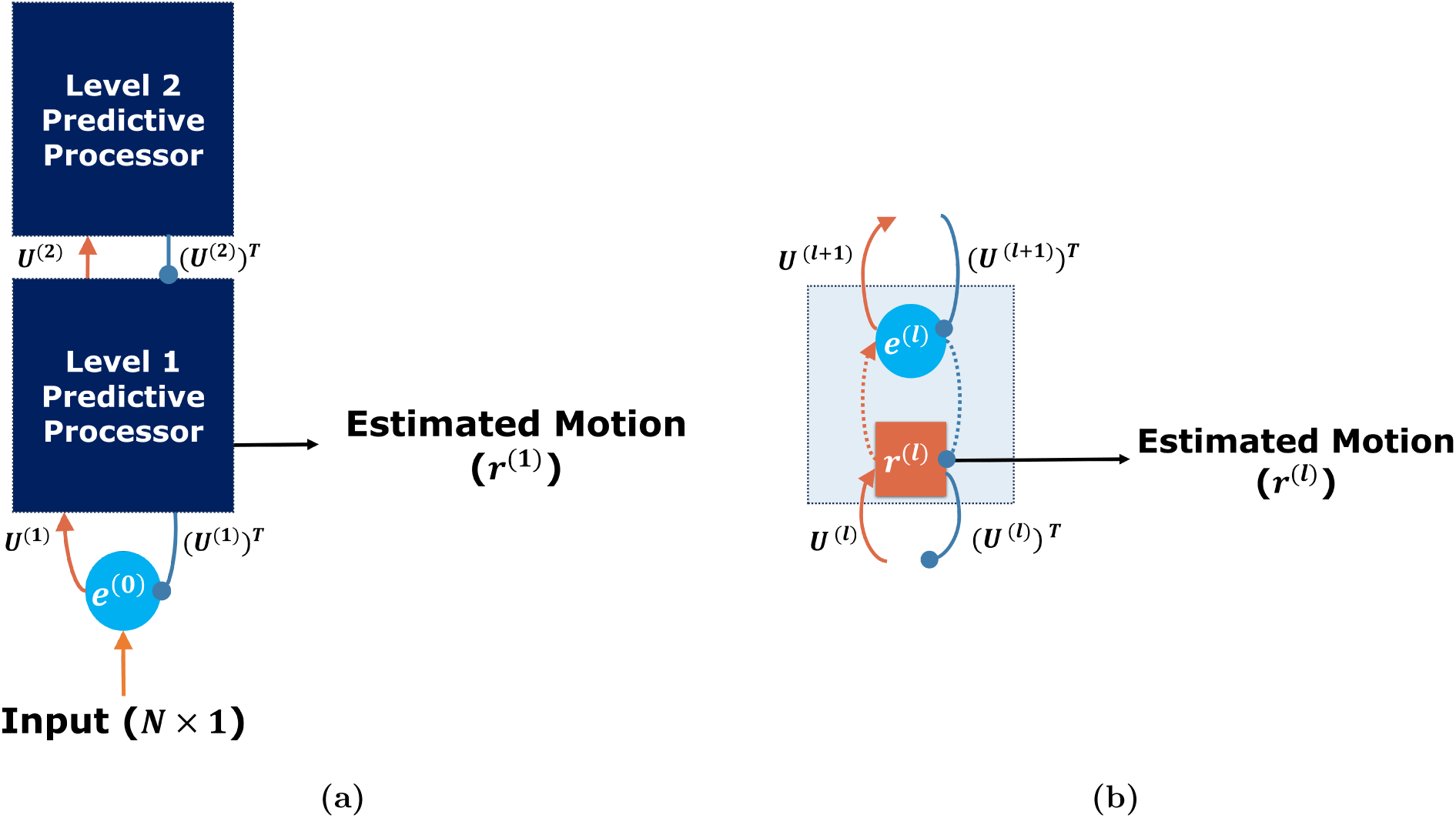
Classical Predictive Coding for motion processing. (a) Two-level classical predictive coding. (b) One level of a predictive processor. Both subplots illustrate the network’s outputs with black arrows. Excitatory and inhibitory connections are highlighted in orange and blue lines, respectively. Error units are depicted as blue circles, while activity units are shown as orange rectangles. Synaptic weights, marked with ‘**U**^(*l*)^’, are displayed on the arrows, where *l* indicates the layer in the hierarchy. Dotted orange and blue arrows represent internal excitatory and inhibitory connections. Here, *l* is the level index, and *N* is the number of input objects. Adapted from Rao and Ballard (1999).

The predictive coding model centers on two types of units within each level: error units and activity units, represented by a blue circle and an orange rectangle in Figure. 3b, respectively. Error units identify mismatches in the brain by comparing predictions against sensory inputs to pinpoint discrepancies, termed “prediction errors,” and relay these errors to subsequent levels in the hierarchy. Activity units create predictions from sensory data and previous prediction errors, transmitting these predictions to the preceding level in the hierarchy.

The predictive coding model aims to minimize the prediction error at each level of the hierarchy within the network. The model achieves this by implementing gradient descent on the prediction errors to adjust connection strengths within the network accordingly. The objective function, *E*, represents a sum-of-squares error:

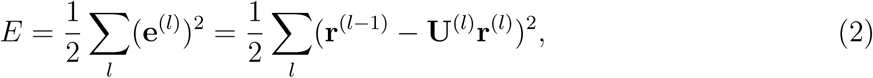

where **e**^(^*^l^*^)^ represents the prediction error at level *l*, **r**^(^*^l^*^)^ is the output of the activity units, denoted as the prediction, and **U**^(^*^l^*^)^ specifies the matrix of synaptic weights that determine the prediction strength. The model treats **r**^(0)^ as the direct sensory input and iterates through levels to minimize *E*.

Consequently, The predictions **r**^(*l*)^ at each level *l* update over time according to the following equation:

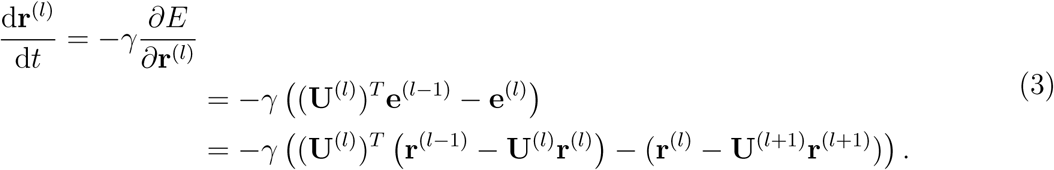

Each level *l* updates its prediction, **r**^(^*^l^*^)^, based on two factors: (1) the feed-forward input as the prediction error, **e**^(^*^l^*^−1)^, from the preceding level, adjusted by the synaptic weight, **U**^(^*^l^*^)^, represented by the solid orange line in Figure. 3b, and (2) the difference between the current prediction, **r**^(^*^l^*^)^, and the feedback from higher levels, **U**^(^*^l^*^+1)^**r**^(^*^l^*^+1)^, illustrated by the dotted blue line in Figure. 3b. The update rate, *γ*, determines the extent to which these inputs influence the update of the current prediction, **r**^(^*^l^*^)^, ensuring a balance between incorporating new data and preserving prior predictions. This equation methodically enhances predictions by applying both feedback and the propagation of errors and adjustments, highlighting the model’s adaptive learning capabilities.

The model updates the synaptic weights to reflect the learning process using gradient descent, expressed by the following equation:

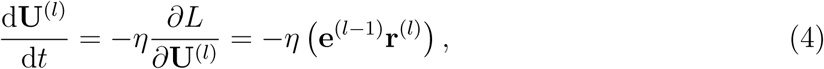

where *η* represents the learning rate, this equation applies the Hebbian rule (Hebb, 1949), demonstrating that the weight update is proportional to the product of the post-synaptic activity, **r**^(^*^l^*^)^, and the pre-synaptic activity, **e**^(^*^l^*^−1)^. The model iteratively adjusts the weights until it converges to an optimal set of weights, indicated by a minimum prediction error.

The current study employs a two-level classical predictive coding algorithm. The model updates the dynamics of the activity unit in predictive coding using the following equations:

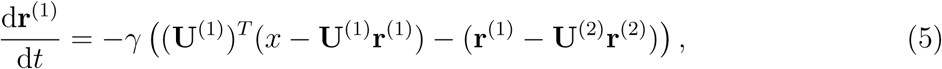

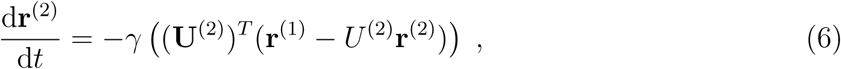

where **r**^(^*^l^*^)^ represents the output of the activity unit, and *x* denotes the input data, specifically the positions of objects within the environment. The variable **U**^(^*^l^*^)^ specifies the synaptic weight that modulates the influence of activity units across the network levels, with subscripts 1 and 2 indicating the first and second level indices, respectively. This hierarchical update process integrates information across levels: the update of **r**^(1)^ depends on the current estimates of **r**^(2)^ along with the synaptic weights **U**^(1)^ and **U**^(2)^. Similarly, the update of **r**^(2)^ incorporates the current estimates of **r**^(1)^, **U**^(1)^, and **U**^(2)^. This iterative interaction ensures dynamic adjustments within the network, enhancing predictive accuracy at both levels.

The model assumes that synaptic weight updates occur on a slower time scale than the rapid updates of neural activities. This assumption reflects the physiological understanding that synaptic plasticity modifies synaptic weights over extended periods and drives long-term learning processes. In contrast, neural activity involves neurons firing in response to stimuli and operates on a much shorter time scale, enabling rapid information processing on a trial-by-trial basis.

To draw a parallel with kinematics, consider the dynamic equation 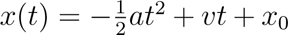, which relates an object’s position *x*(*t*) over time *t* to its acceleration *a* and velocity *v*, with *x*_0_ as the initial position. Our model aligns **r**^(2)^, representing an estimate of the system’s acceleration, with the acceleration term *a*, and matches **r**^(1)^, representing an estimate of velocity, to the velocity term *v*. This interpretation emerges because lower-level representations like **r**^(1)^ respond to immediate changes in input data, similar to velocity, while higher-level representations like **r**^(2)^ capture slower, broader adjustments, akin to acceleration in kinematics.

The analogy highlights the hierarchical nature of the model, with **r**^(1)^ (velocity) responding directly to input data changes and **r**^(2)^ (acceleration) representing generalized, slower-changing features of dynamics. Our main assumption posits that the motion of each object results from the weighted sum of its motion sources, interpreting each motion source as representing velocities. The model leverages the predictive coding framework to represent each motion source, where **r**^(1)^ in each predictive coding unit corresponds to the motion source, and the synaptic weight **U**^(1)^ quantifies the influence of each motion source on the object.

At each time step, the model forwards the estimated motion derived from the predictive coding framework to the surround suppression mechanism for refinement. The following section explains the surround suppression mechanism’s operation within the predictive coding framework.

### 2.3 Surround Suppression Mechanism

The proposed model places surround suppression at its core, as illustrated in Figure. 4. This mechanism modulates the response of motion-sensitive neurons to a stimulus within the central receptive field based on concurrent motion stimuli in the surrounding areas, aligning with principles outlined by Born (2000).

**Figure 4:**
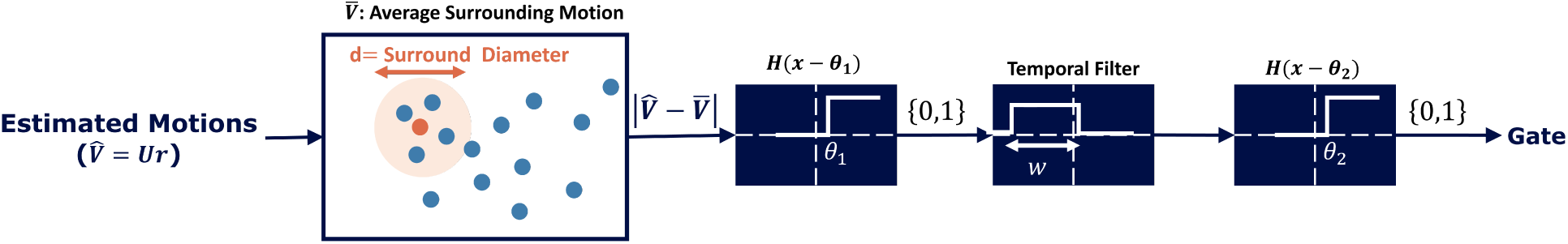
Surround Suppression mechanism. This diagram compares motion estimation at the center of the receptive field, denoted by *v̂* = *Ur*, with the average motion in the surrounding area, denoted by *V̄*, at each temporal step. This comparison uses a Heaviside function with the threshold parameter *θ*_1_. The result is then integrated with the historical average motion within a temporal width window *w*. This integrated value is again subjected to comparison, passing through a Heaviside function with a threshold parameter *θ*_2_. The output is binary, indicating the activation state of the surround suppression mechanism.

The model links center-surround modulation to extra-classical receptive field effects observed in cortical neurons, particularly in the middle temporal (MT) area, rather than retinal ganglion cells (RGCs), where such interactions are more commonly associated. In cortical neurons, a stimulus in the surround does not directly elicit a response but instead modulates the response to the center stimulus. This modulation works antagonistically for direction-selective neurons: surround motion aligned with the centre’s preferred direction suppresses activity while opposing motion enhances it (Albright, 1984). The model further postulates that the surrounding suppression mechanism’s historical activity also influences this modulation.

The surround suppression mechanism stays inactive when the motion of an individual object aligns with the motion within the surround. The mechanism activates when a misalignment occurs between the surround and the object’s motion estimate. The computation incorporates weighted historical activity to account for temporal dynamics. When activated, the surround suppression mechanism suppresses the neuron’s activity. Conversely, when it remains inactive, the mechanism does not suppress the object’s estimate in the center of the receptive field, allowing the model to distinguish the object’s motion from the surrounding activity.

To implement the surround suppression activation function, the model compares the motion estimation of an object, **v̂** = **Ur**, with the average motion observed within the object’s surroundings, which the model assumes to be circular with a diameter of *d*. The model defines this average motion as

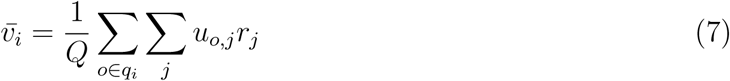

where *q_i_* defines the set of indices corresponding to objects within the circular area surrounding object *i*, and *Q* represents the total number of these surrounding objects. The double summation computes the combined influence of all motion sources (indexed by *j*) on each surrounding object *o*, estimating their average motion in the surround.

In our model, the surround suppression mechanism accounts for motion within a specific region around the center of the receptive field. This method treats all surrounding objects as contributing equally to the calculation, regardless of their spatial distance from the receptive field center. While this equal-weight approach ensures computational efficiency, it simplifies the model by not accounting for distance-dependent synaptic variability. However, we did not enforce specific patterns on the shapes of synaptic activity based purely on distance. Instead, the synaptic weights are shaped through a learning process that reflects how neurons respond to motion over time, integrating the influence of surround motion in a biologically plausible manner. A distance-weighted averaging scheme in future studies could improve the model’s biological realism by accurately representing synaptic interactions.

The surround suppression module governs its behavior by actively integrating current and historical motion data through the function:

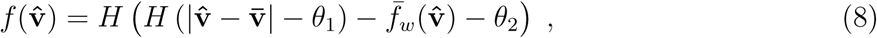

where the Heaviside step function *H*() determines the activation state. The module evaluates the motion similarity by comparing the individual object’s motion, *v̂_i_*, with the average motion of its surrounding objects, *v̄_i_*, using the threshold *θ*_1_. It further incorporates historical influence by applying the secondary threshold *θ*_2_, which modulates the current response based on the historical average of suppression activation. The module uses these thresholds to actively assess and adjust the surround suppression mechanism, balancing immediate and historical factors to refine its output.

The parameter *θ*_1_ plays a pivotal role by setting the threshold for surround suppression, determining how the model compares the motion within the center of the receptive field to the collective motion in the surrounding visual field. Increasing *θ*_1_ raises the threshold for surround suppression, requiring a larger difference between the estimated and average motion in the surround to activate the suppression mechanism. This adjustment directly influences how the model prioritizes or suppresses motion signals based on their divergence from the surrounding context.

The historical average of surround suppression activation, *f̄_w_*, is calculated over a specified time window *w*. This temporal filter integrates information from past suppression activations, ensuring the model captures temporal dynamics in motion prediction. By broadening the window *w*, the model incorporates a more extensive history of motion data, reducing the influence of transient noise and enhancing the stability of coherent motion detection. The “window duration” refers to the size of the temporal window used in the temporal filter, as expressed in (8). This window size determines how much past information from the surround suppression mechanism is considered during motion detection. The temporal filter integrates information over this window to allow the model to account for prior motion activity in the surrounding regions. Here, noise reflects variability in external sensory input, such as random dot motion, and its impact on simulated neuronal responses. The model excludes internal neural noise, such as variability in firing rates or synaptic activity, to simplify the dynamics and focus on the core processes being studied..

This study examines how the parameters *θ*_1_ and *w* affect the model’s ability to detect coherent motion. The model uses *θ*_1_ to process immediate motion within its receptive field and relies on *w* to define the duration of the temporal window for evaluation. While *θ*_2_ governs the integration of past motion information and influences how the model distinguishes relevant from irrelevant historical movements, this study does not focus directly on *θ*_2_. Instead, it indirectly addresses its effects through the temporal window duration *w*. Our primary objective is to assess how incorporating historical motion data improves the model’s capacity to identify coherent motion in real-time sensory input. By fine-tuning the balance between immediate sensory cues and accumulated historical data, we aim to enhance the model’s robustness and accuracy in detecting coherent motion patterns.

## 3 Mechanistic Framework of the Model

The proposed model is grounded in a two-level predictive coding framework enhanced by surround suppression, designed to iteratively refine motion estimates. Predictive coding is the core computational strategy, integrating sensory inputs with prior knowledge to facilitate motion detection. Surround suppression introduces two key parameters—windows and thresholds—that shape this process. Inspired by neural mechanisms, these parameters, inspired by neural mechanisms, reflect biological processes such as synaptic integration and inhibition. The windows of integration accumulate historical motion data over a temporal span, mimicking how neural circuits process inputs over time. Thresholds define the minimum motion contrast between the center and the surround required to activate suppression, capturing spatial and temporal modulation of neuronal excitability.

Temporal integration further enhances the model by incorporating the historical activation of surround suppression within a specific time window, creating a robust understanding of motion patterns over time. If suppression was previously activated during this period, it influences current neuronal activity, simulating the impact of prior information on motion detection. The model effectively simulates real-time motion detection while considering historical visual inputs by combining spatial integration through vector analysis with temporal integration through experience binding.

The hierarchical framework distinguishes between individual and shared motion. Individual motion arises from the first level of predictive coding, assessing the local motion of objects based on direct sensory input. In contrast, shared motion emerges after applying surround suppression, which integrates neighboring motion cues to identify global patterns. The model does more than simply reproduce the well-known fact that higher-level representations tend to become more global. The contribution of this model lies in the ability to simulate the intricate balance between local motion detection and global motion integration by incorporating both predictive coding and surround suppression. This nuanced approach allows for the decomposition of motion structure, as demonstrated in experiments such as the Random Dot Kinematogram (RDK), where shared motion emerges from noise-dominated environments, and the complex structure of the Johansson and Dunker wheels experiments. The model advances our understanding by quantitatively demonstrating how this interplay between local and global motion components evolves over time, especially in relation to motion perception deficiencies observed in schizophrenia.

Surround suppression is crucial in identifying coherent motion patterns while suppressing noise or conflicting inputs. The model computes the prediction error at each stage—the discrepancy between the estimated motion and actual sensory input—which is propagated to the next stage for iterative refinement. The surround suppression mechanism specifically targets the detection of global motion in stimuli. By comparing motion signals between the center and the surrounding areas of the receptive field, the model activates suppression to extract shared motion in the visual scene. This spatial analysis, represented as vector comparisons, detects deviations or consistency in motion patterns, forming the foundation for identifying coherent motion signals.

The model incorporates synaptic weight modulation across two timescales to simulate neural adaptability. SThe short-term modulation captures immediate changes in synaptic strength in response to stimuli, although it is not as fast as the real-time changes in neural activity. This allows our model to reflect how synaptic weights can adjust over the course of a single trial. Additionally, we have incorporated long-term synaptic plasticity by relying on past information to adjust the synaptic weights over extended periods. This dual approach allows the model to simulate both rapid responses to new sensory input and the gradual influence of prior experiences on motion detection. We recognize that modeling the interactions among motion sources using synaptic weights is an abstraction and does not fully capture the distinct physical and temporal differences between these processes. The correspondence between synaptic weight changes and neural activity modulation in our model is intended to be instrumental, aiding in exploring noise’s role in motion detection and related deficiencies.

The hierarchical structure of the model reflects the brain’s ability to process global and local motion simultaneously. The initial module prioritizes global motion to establish a contextual foundation, while subsequent modules refine shared motion representations and resolve individual motion. This structure captures the dynamic interaction between global contextual processing and local precision. By abstracting motion interactions through synaptic weight changes, the model provides a functional framework for studying the impact of noise on motion detection and understanding hierarchical neural processing in dynamic environments. The combination of spatial and temporal integration ensures the model effectively simulates real-time motion detection while incorporating prior experiences.

### 3.1 Stimuli

We evaluated the algorithm using simulation results from three psychophysical experiments: (I) the Johansson experiment (Johansson, 1973), (II) the Duncker wheel (Duncker, 1929), and (III) the Random Dot Kinematogram (Kim and Shadlen, 1999). Each experiment contributes uniquely to the study of motion perception and processing.

In the Johansson experiment (experiment I) (Johansson, 1973), illustrated in Figure. 5a-b, three vertically aligned dots move back and forth across a display. The two outer dots move horizontally, while the middle dot moves diagonally, maintaining collinearity with the outer dots (Figure. 5a). Observers perceive the middle dot as moving vertically, oscillating between the two horizontally moving outer dots, despite its actual diagonal trajectory (Figure. 5b). This illusion arises from the synchronized horizontal motion of all three dots, where the brain interprets movement based on their relative positions and synchronized movements.

**Figure 5:**
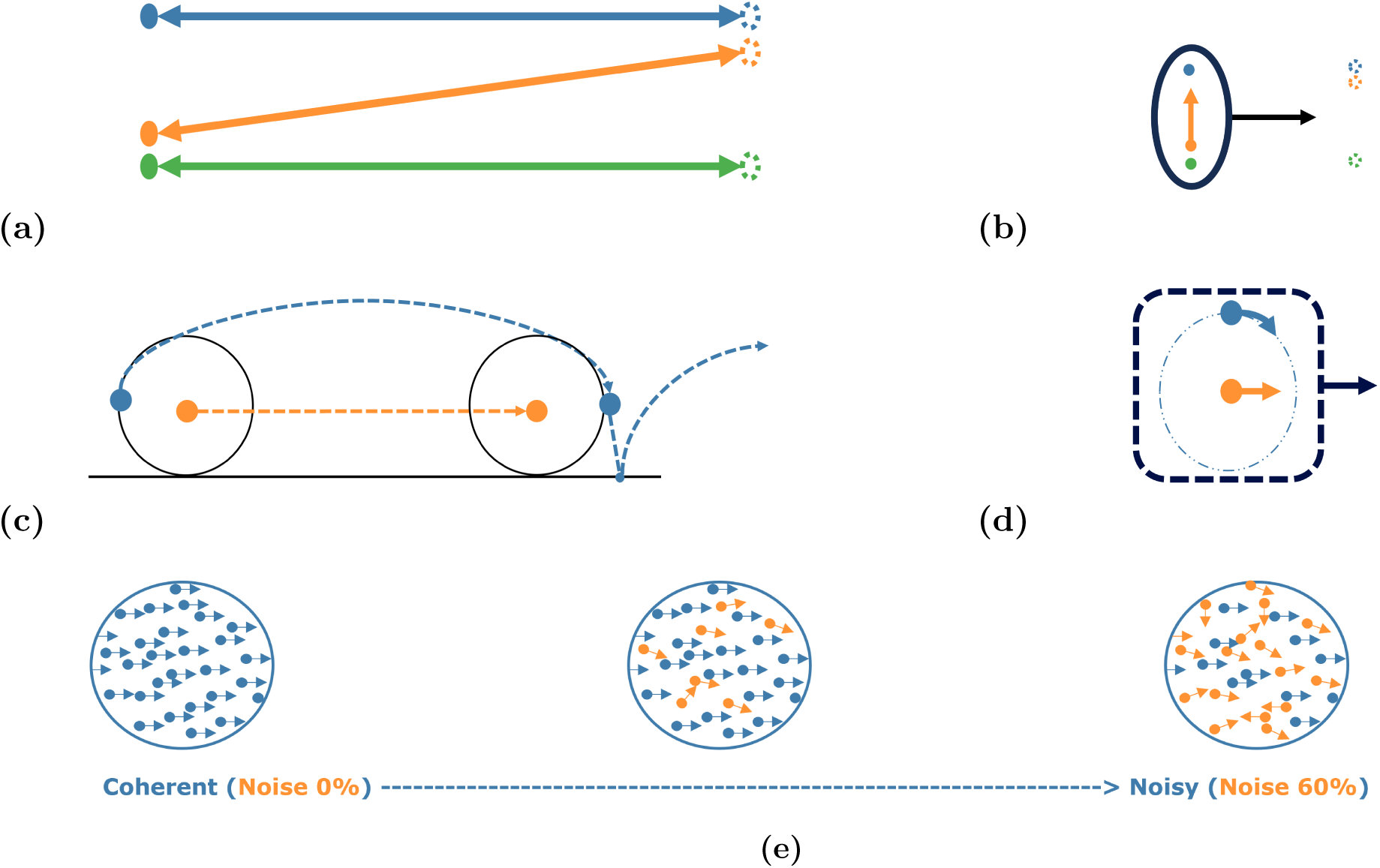
Experimental Paradigms Overview. (a-b) A schematic view of Johansson’s experiment with three dots, showing (a) motion vectors for each of the dots and (b) the perceived motion of the group (denoted by the oval containing the three dots moving rightwards) and the central dot (moving upwards). Adapted from Gershman et al. (2016). (c-d) A schematic view of Duncker wheel’s experiment with three dots showing (c) motion vectors for each of the huband rim-dots, and (d) the perceived motion of the group (denoted by the dashed square containing both dots moving rightwards) and the rim-dot (moving in a circular motion). Adapted from Gershman et al. (2016). (e) Random Dot Kinematogram. The difficulty of the experiment escalates from left to right with an increasing number of noise dots, depicted in orange (although the dot colors are not different in the experiment). The blue dots represent signal dots that have coherent motion towards the right.

The Duncker wheel experiment (experiment II) (Duncker, 1929) explores motion perception by featuring two dots placed on an invisible wheel, as shown in Figure. 5c-d. One dot, positioned at the hub of the wheel, moves linearly (typically left to right), while the other dot follows a complex spiral path along the rim due to the wheel’s rotational and lateral movement (Figure. 5c). Observers perceive these motions differently: they see the hub dot moving linearly, consistent with its actual motion, but perceive the rim dot, despite its spiral path, as moving in a simple circular motion around the hub dot (Figure. 5d). This phenomenon demonstrates the brain’s tendency to break down complex movements into recognizable components, interpreting the spiral motion of the rim dot as a combination of translation and rotation when the hub dot serves as a reference point.

The Random Dot Kinematogram (RDK) (experiment III) serves as a valuable method for investigating motion perception, particularly in the context of schizophrenia (Kim and Shadlen, 1999). In an RDK, numerous small dots move randomly across a screen, as shown in Figure. 5e. Among these, a subset of dots (blue) moves coherently in the same direction, while the remaining dots (orange) move randomly. Observers are tasked with identifying the direction of coherent motion. For example, in a display with 95% coherence, 95% of the dots (signal dots) move in a designated direction between frames, while the remaining 5% (noise dots) move randomly. Higher coherence levels facilitate the perception of global motion direction. This experimental design provides critical insights into how the visual system processes and differentiates motion information from noise. The concept of “time to convergence” is introduced to evaluate model performance in this context. This refers to the number of discrete time steps, each with a Δ*t* duration, required for the model to identify the direction of coherent motion within the RDK accurately. Coherent motion is considered accurately identified when the motion strength stabilizes, with variations remaining below 0.05 units across successive time steps. This ensures the model converges to a stable and reliable representation of coherent motion direction.

The computational time (a surrogate measure of latencies) measured in the model does not correspond directly to the time required for cognitive processing in the brain. Instead, the model provides an abstraction of the underlying mechanisms involved in motion detection rather than replicating the exact timing of neural responses. In biological systems, multiple brain regions, including attention mechanisms and higher-order decision-making, contribute to motion perception and influence response times. Accurately modeling these temporal dynamics would require a more detailed framework that includes the complexities of cortical interactions, synaptic plasticity, and attentional modulation, which lies outside the scope of this work.

The parameters of the stimuli used in the experiments are summarized in Table 1. The subsequent section presents the results of the model simulations and their role in elucidating key aspects of human visual motion perception as explored through these psychophysical experiments.

**Table 1:** Parameters of stimuli used in the study.

| Stimulus | Parameter | Value |
| --- | --- | --- |
| Johansson Experiment | Frequency | 0.5 Hz |
| | Horizontal Amplitude | $2\sqrt{0.3}$ |
| | Vertical Amplitude | $\cos(45^\circ)2\sqrt{0.3}$ |
|  | Time Constant | 1/60 s |
| Dunker Wheel | Wheel Radius (R) | 10 unit |
|  | Horizontal Speed | 0.5 unit/s |
|  | Angular Speed | 0.1 deg/s |
|  | Time Constant | 0.05 s |
| Random Dot Kinematogram | Number of Dots | 1000 |
|  | Dot Density | 500 dot/unit |
|  | Coherent Velocity | 0.01 unit/s (horizontal motion) |
|  | Random Velocity | 0.01 unit/s (vertical motion) |
|  | Time Constant | 0.05 s |

## 4 Results

We evaluated the model’s performance in identifying motion structures and extracting coherent motion across three distinct experiments. Furthermore, we analyzed how prior knowledge affects the model’s performance with motion data and examined the role of the surround suppression mechanism in detecting coherent motion.

### 4.1 The model successfully identified the motion structure in classical motion patterns

Figure. 6 illustrates the results of classical motion perception experiments simulated using our predictive coding framework. The model successfully captured the motion structures in the Johansson (1973) and Duncker (1929) experiments (I and II), effectively distinguishing between shared and individual motion types.

**Figure 6:**
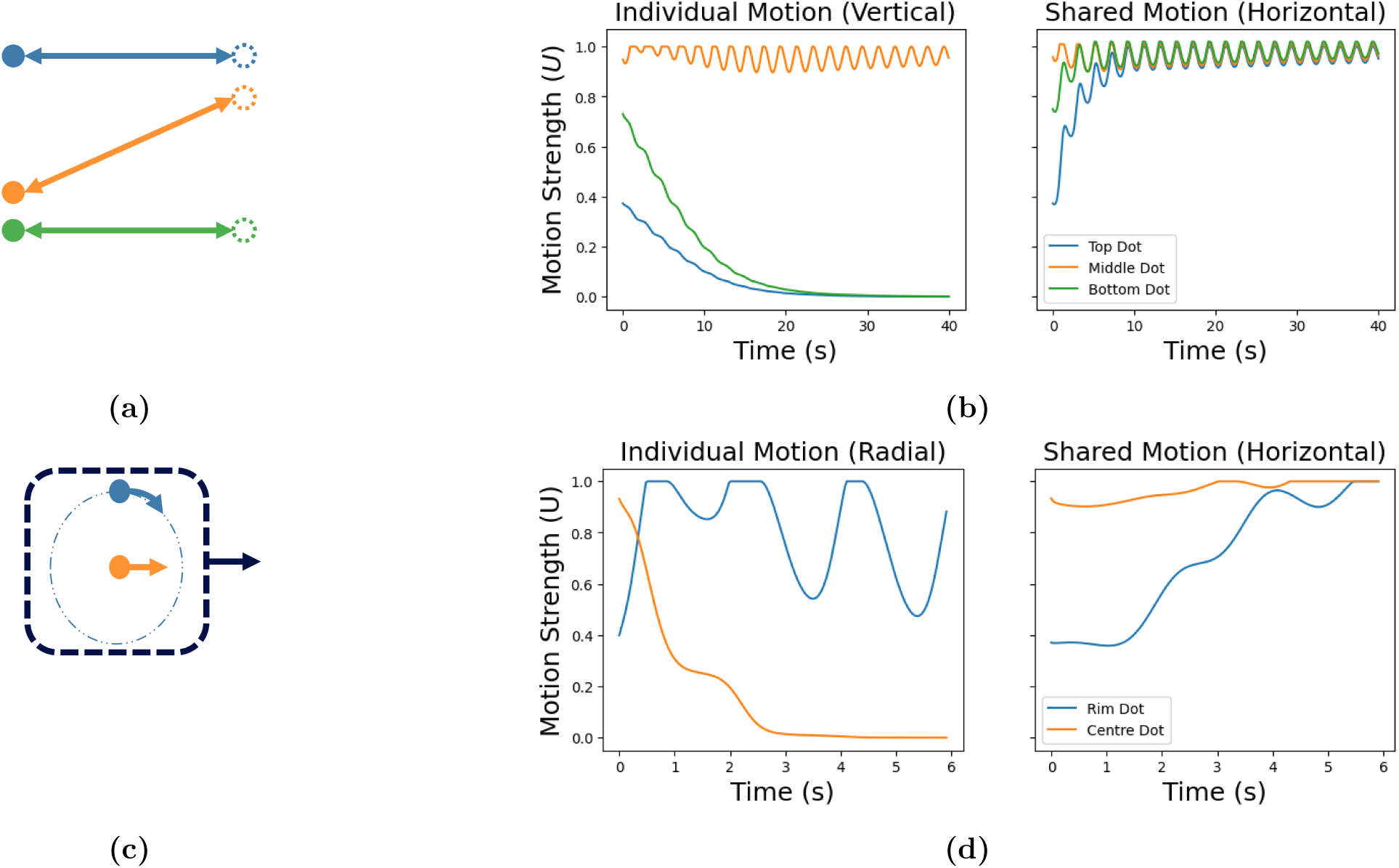
The model successfully identified the motion structure in experiments I and II. (a) A schematic view of Johansson’s experiment I (Johansson, 1973) with three dots. (b) Model estimates of motion strength: shared and individual, horizontal and vertical motions, respectively. The individual motion strengths reach zero for the two horizontal moving dots (blue and green), but is close to maximum for the middle dot (orange).(c) The Duncker wheel experiment II (Duncker, 1929) with a central dot (orange) and a dot on a circle (blue). (d) The model recognises shared motion (horizontal motion) and an individual element (rotational motion) of the rotating dot. The plots are colour coded, matching the colours of the objects with their corresponding plot lines.

Psychophysical experiments on Johansson motion in experiment I (see Figure. 6a) show that humans perceive this stimulus as shared horizontal motion, with the central dot oscillating vertically (Johansson, 1973). Based on these observations, we expect the model to extract equal (synchronous) shared motion across all three dots and to identify individual motion specifically for the central dot. The model’s output consists of connectivity weights **U**, which represent the strength of the contribution of each motion source (motion strength). Figure. 6b displays the estimated motion strengths for shared and individual motions. The individual motions of the blue and green dots converge to zero over time, while the individual motion of the central dot (orange) remains pronounced. The shared motion of all three objects converges to similar values, close to 1. These results demonstrate the model’s ability to distinguish between shared motion, where all dots synchronize, and individual motion, which diminishes for the outer dots but persists for the central dot. The observed oscillation in motion strength arises from the sinusoidal motion of the objects as they move back and forth horizontally. In this experiment, shared motion corresponds to the collective horizontal movement of all dots, causing synchronized oscillation. Individual motion is characterized by vertical movement, which is absent for the blue and green dots (vertical motion strength is zero) but present for the orange dot, as indicated by its non-zero motion strength.

The application of the model to the Duncker experiment in Experiment II (Figure. 6c) validates its ability to represent complex motion patterns. As illustrated in Figure. 6d, the model successfully captured the shared horizontal motion of the dots while distinguishing the individual motion of the rim dot (blue) through its significant individual motion strength. In contrast, the other horizontally moving dot (orange) exhibited an individual motion strength that converged to zero, effectively simulating the perception of a rolling wheel. In this experiment, shared motion corresponds to the horizontal movement of both dots, causing them to move horizontally to the left. Individual motion, interpreted as radial motion, converges to zero for the horizontally moving dot but remains non-zero for the rim dot (blue). The pronounced oscillation observed in the shared motion of the radial ball, resulting from the consistent value of the motion source, effectively represents the motion strength of the radial movements of the blue dot. The period and amplitude of the sinusoidal motion strength output vary with several experimental variables. For example, changes in the positions of dots and the duration of their movement directly influence the amplitude and frequency of the oscillation. When the objects follow a more structured pattern or move over a longer duration, the model produces a more stable oscillation with a higher amplitude, accurately reflecting the motion’s dynamics.

Figure. 7 presents the results from a Random Dot Kinematogram (RDK) in experiment III, which illustrates the distinction between individual and shared motion. Noise dots (dots with noncoherent motion) exhibit higher and more variable individual motion strengths compared to signal dots (orange line). Despite this, the large number of noise dots prevents their individual motion strength from reaching 1. The mean individual motion strength for noise dots ( 10.4 units/ms) exceeds that of signal dots ( 3.25 units/ms), with noise dots displaying significant fluctuations while signal dots maintain a relatively stable, lower value. This contrast underscores the unpredictable and incoherent movements of noise dots compared to the coherent motion of signal dots. In the shared motion graph, signal dots quickly reach and sustain a high shared motion strength close to 1, whereas noise dots exhibit almost no shared motion. This indicates that the model successfully isolates the coherent motion from the random background. As the model stabilizes, the individual motion strengths of signal dots diminish to zero, demonstrating the successful detection of coherent motion. The drop in motion strength for the signal dots at approximately 100 ms likely marks the point at which the model begins to recognize not only the shared motion but also the individual noisy motion of the dots. Up to 100 ms, the model primarily reinforces the presence of shared motion in the stimuli. Beyond this point, the model incorporates the contribution of individual dot motion, explaining the observed decline in overall motion strength. This dynamic highlights the model’s ability to adjust and integrate motion information over time.

**Figure 7:**
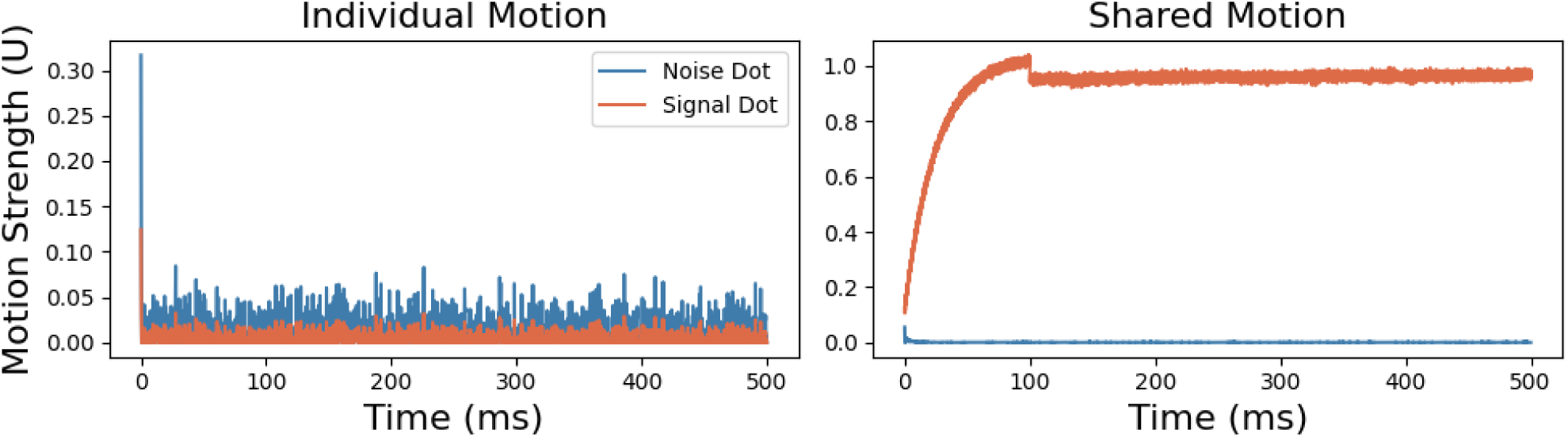
Individual and shared motion in a random dot kinematogram, experiment III. Blue lines represent the motion strengths of noise dots, while the orange lines depict the motion strengths of signal dots with coherent motion. Note the reduced scale axis of the individual motion plot, used to highlight the low motion strengths of the dots.

### 4.2 The Balance between sensory data and prior knowledge in coherent motion detection

The model’s performance in detecting coherent motion in the RDK task was assessed by measuring the time to convergence. Figure. 8 illustrates variations in convergence time as noise levels in the stimuli change, along with the impact of prior knowledge on the model’s performance. Figure. 8a shows a strong relationship between the signal-to-noise dots ratio and convergence time, with higher noise levels leading to longer convergence durations. This experiment was conducted for historical window durations of 0, 10, 30, 50, 70, and 90 ms, with 150 trials per window duration.

**Figure 8:**
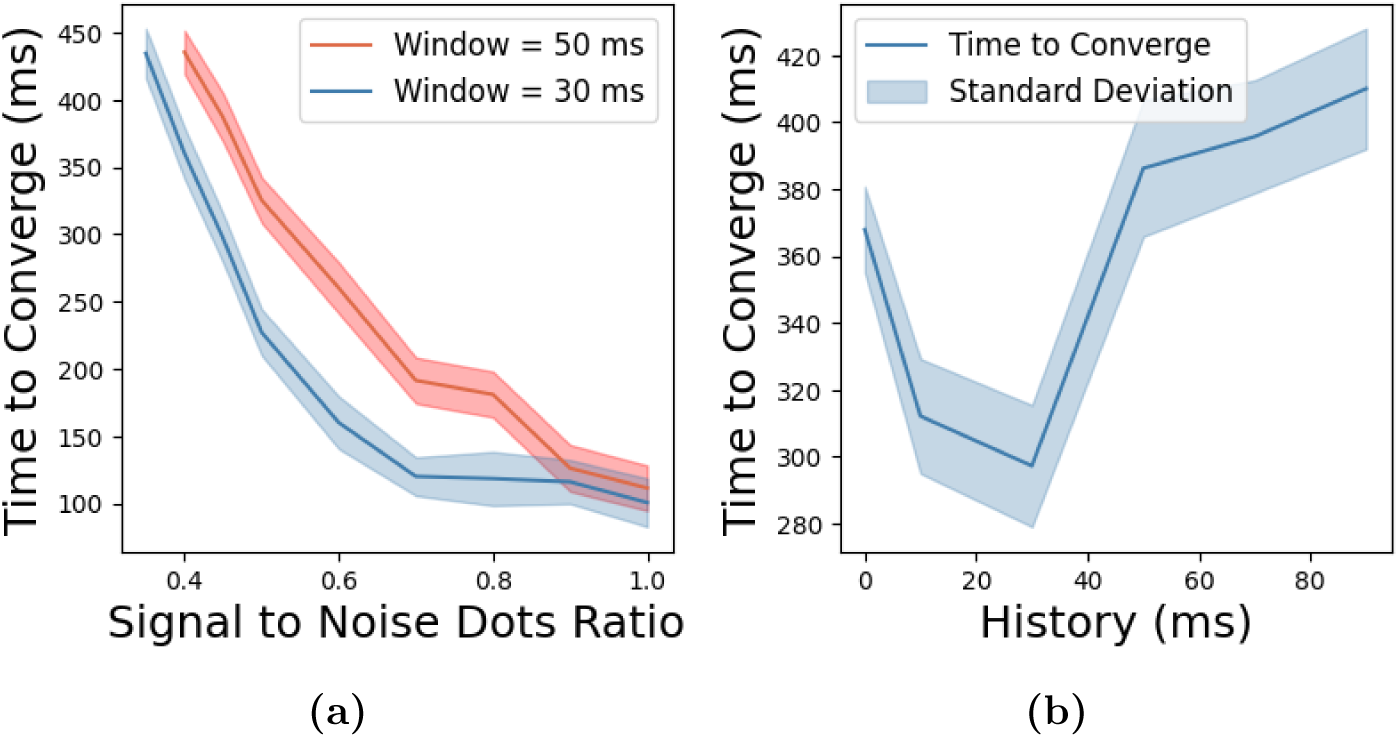
Effects of noise and historical window duration on the time to convergence of model (experiment III). (a) The time to converge as a function of signal to noise dots ratio for two different historical window durations: 50 ms and 30 ms.(b) Convergence time’s dependence on the historical window duration. The shaded areas around the lines represent the standard deviations across 150 trials for each window duration.

Figure. 8a compares the effect of signal-to-noise dots ratio on convergence time for two historical window durations: 50 ms and 30 ms. The longer historical window resulted in longer convergence times compared to the shorter window. The shaded regions in the figure indicate the standard deviation across the 150 trials.

Figure. 8b depicts the relationship between historical window duration and convergence time for a signal-to-noise dots ratio of 0.4. Convergence time initially decreases as the historical window duration increases, reaching an optimal point at 30 ms. Beyond this point, further increases in historical window duration lead to a rise in convergence time. These findings suggest an optimal range of historical information that enhances the detection of coherent motion. Exceeding this range appears to hinder the model’s performance, likely due to an over-reliance on historical data.

In Figure. 8, we present results for window durations of 30 ms and 50 ms, as these durations demonstrate the model’s performance across varying noise proportions. We chose to highlight the 30 ms window because it achieves an effective balance between integrating past information and maintaining a computationally efficient time to convergence. As shown in Figure. 8b, the time to converge increases with higher noise proportions, yet the model exhibits a consistent trend across different noise levels. The results for the 30 ms and 50 ms windows demonstrate stability under varying noise conditions, indicating the robustness of the model. These findings suggest that the model effectively adapts to different noise levels while maintaining reliable performance across these historical window durations.

### 4.3 Surround suppression and the detection of coherent motion

The model’s performance was assessed by modifying the influence of surround suppression on the detection of coherent motion in the RDK task (experiment III). Specifically, we investigated the effect of adjusting the threshold parameter (*θ*_1_) in the surround suppression mechanism. A higher threshold value indicates that the motion in the center and surround must differ significantly for the surround suppression mechanism to activate.

Figure. 9 summarizes the model’s performance in the RDK task. The axes represent the surround suppression threshold and the signal-to-noise dots ratio. The color gradient reflects the convergence time, with cooler colors indicating shorter durations and warmer colors indicating longer durations. The figure demonstrates that as noise levels increase, the time to converge also increases, indicating that the model requires more time to identify coherent motion directions under higher noise conditions.

**Figure 9:**
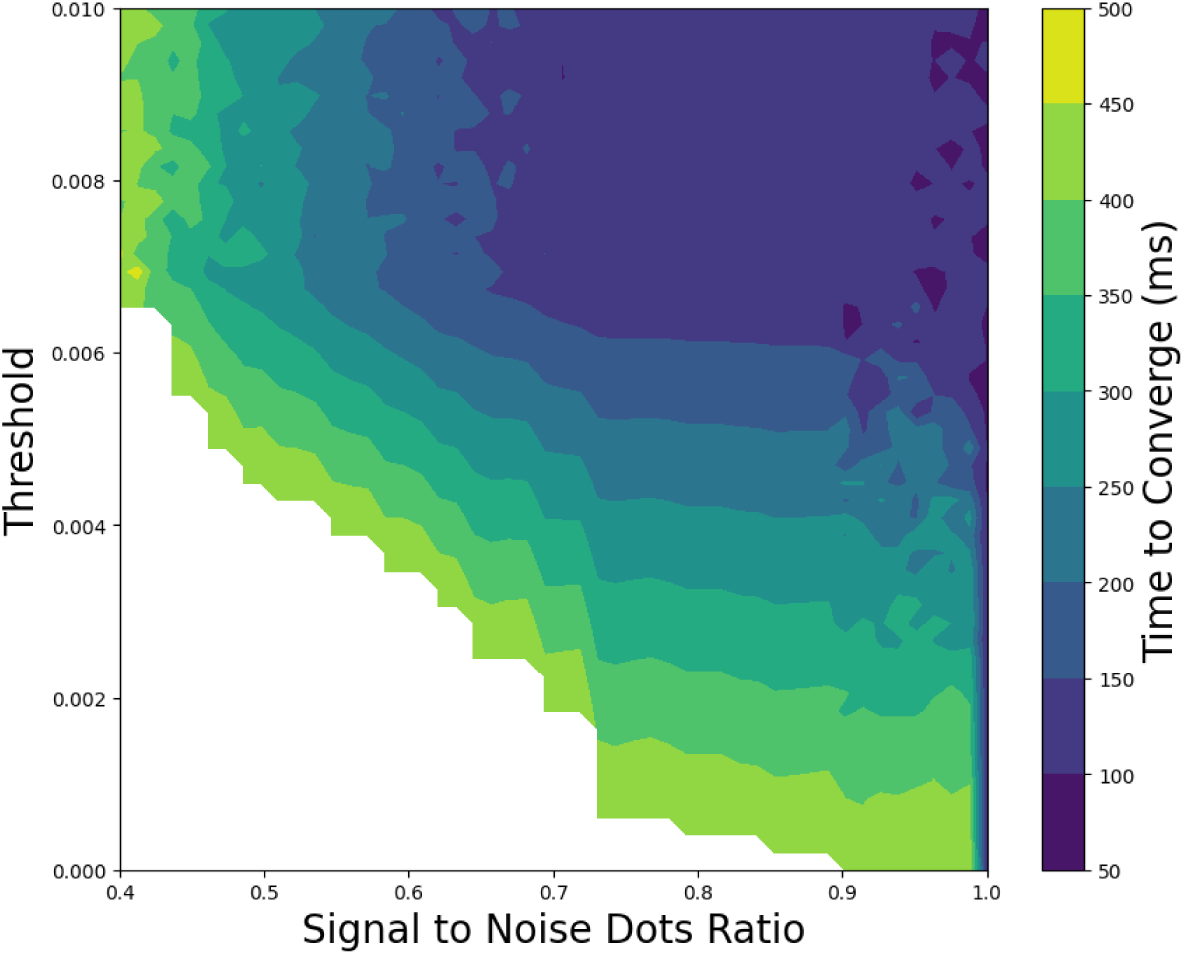
Effects of noise level and surround suppression threshold on model convergence time in the RDK task, experiment III. The color indicates the convergence time of the model in ms, with cooler colors representing shorter times and warmer colors indicating longer times.

The results also highlight the impact of the surround suppression threshold on convergence time. Lower thresholds result in longer convergence times, while increasing the threshold to sufficiently high levels enhances the model’s performance in noisy environments. Higher thresholds reduce the impact of noise on the surround suppression mechanism, preventing excessive changes due to random noise and improving the model’s ability to process coherent motion.

## 5 Discussion

This study presents a biologically plausible computational model based on the predictive coding algorithm for coherent motion detection. The model extracts motion structure from the visual field by decomposing motion into individual and shared sources, providing insights into motion perception and its potential deficiencies in schizophrenia. It integrates core processes such as predictive coding and surround suppression, fundamental mechanisms that link neural processes to behavioral outcomes. Although the model adopts an abstract approach without explicitly simulating specific neurons or cortical layers, it offers a broad and adaptable framework for understanding motion detection.

Center-surround interactions in the middle temporal (MT) area are critical for motion perception, as the activity of direction-selective neurons is modulated by stimulus alignment with its surround through suppression or enhancement (Allman et al., 1985; Bradley and Andersen, 1998; Tadin et al., 2003; Eskikand et al., 2018;Zarei Eskikand et al., 2019). These mechanisms contribute significantly to coherent motion perception (Albright, 1984).

Our model successfully replicates key findings from psychophysical experiments, particularly in extracting motion structures observed in the seminal studies by Johansson (1973) and Duncker (1929). Despite significant advancements in understanding motion perception, few studies have explored this domain in the context of schizophrenia, leaving a critical gap in the literature. The proposed model bridges this gap by linking neural mechanisms to psychophysical outcomes, shedding light on motion perception deficits in schizophrenia. The model replicates findings of weaker surround suppression in schizophrenia (Serrano-Pedraza et al., 2014).) and, for the first time, offers a mechanistic explanation for how motion detection deficiencies observed in some motion detection tasks might arise from this impaired surround suppression.

This finding is crucial as it suggests a new hypothesis that observed weak surround suppression could underlie the motion detection deficiencies observed in patients with schizophrenia. No previous model has explicitly demonstrated this connection, making our contribution both novel and significant. Furthermore, this understanding could pave the way for developing clinical or diagnostic tools to better assess and manage motion perception impairments in schizophrenia, potentially aiding in early detection and personalized interventions.

Additionally, prior research has shown that individuals with schizophrenia often require more prolonged exposure to detect global motion in noisy environments (Spencer et al., 2013). By adjusting the influence of prior knowledge and the strength of surround suppression, our model provides a novel explanation for these deficits, offering valuable insights into the neural basis of motion perception differences between healthy individuals and those with schizophrenia.

The model also proposes that motion perception deficits in schizophrenia may arise from either an over-reliance on prior information (under-learning) or excessive adaptation to new sensory feedback (over-learning). This hypothesis aligns with broader theories suggesting that schizophrenia involves disruptions in balancing sensory input with prior expectations (Kaplan et al., 2016; Stephan et al., 2016; Corlett et al., 2019; Horga and Abi-Dargham, 2019; Nassar et al., 2021). By incorporating these dynamics, the model provides a mechanistic explanation for the behavioral deficits observed in motion perception tasks. To further explore these deficits, we propose an experiment (illustrated in Figure. 10) that investigates the effects of discordant motion in the surround on coherent motion detection. Participants would judge the direction of coherent motion in the center of stimuli, while the angle of difference between the motion of surround elements is systematically varied. Measuring the minimum duration required to detect coherent motion in the center would allow for a comparison between neurotypical individuals and patients with schizophrenia. This experiment could reveal how angular differences in surround motion impact motion detection and provide insights into the neural circuitry underlying surround suppression deficits.

**Figure 10:**
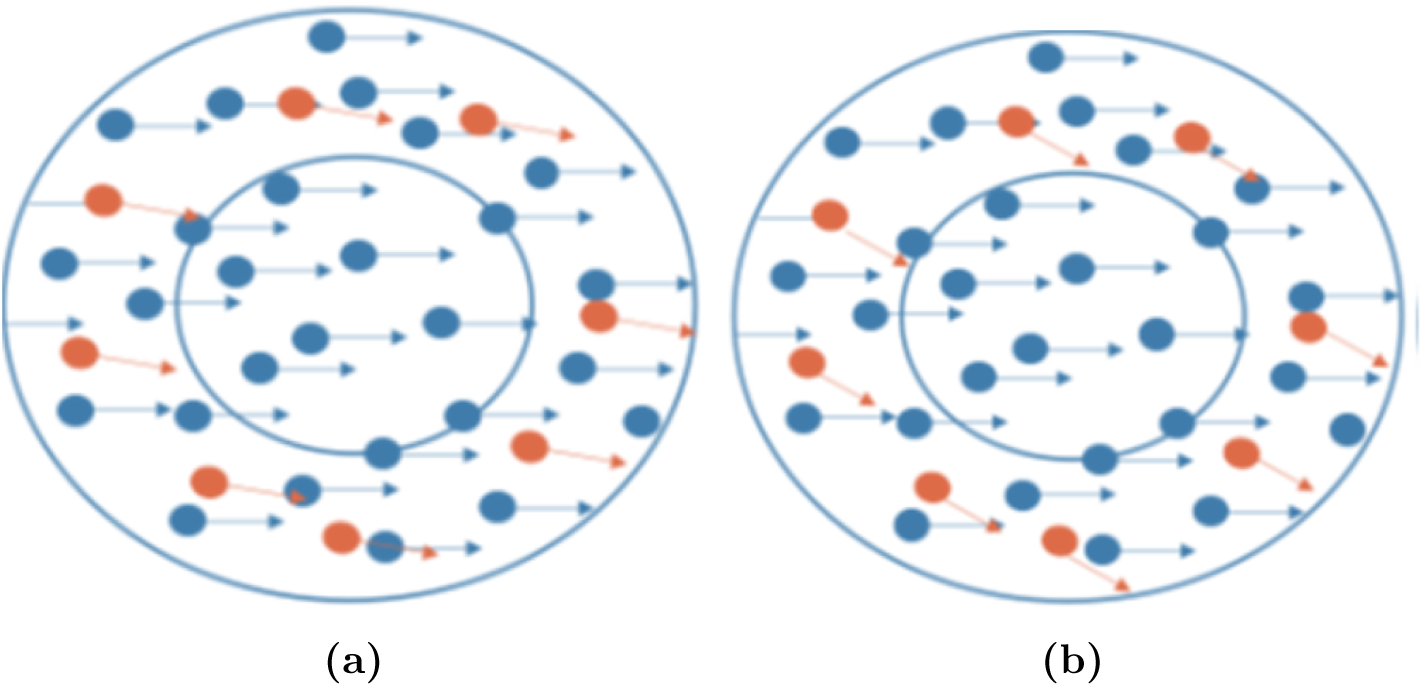
Stimuli to test the coherency threshold for surround suppression. (a) Two sets of overlapping random dots in the center (only blue dots) and the surround (blue and orange dots). Both sets of dots have coherent directions, except that the angle of the motion direction for the red dots is slightly different. (b) The same dots, but the angle of the difference between the motion directions for the red and blue dots is larger.

This study underscores the application of predictive coding in explaining motion perception while laying the groundwork for future research. Expanding the model to incorporate internal noise dynamics, distinctions among cortical areas, and direct links between neural mechanisms and cognitive functions could provide deeper insights into typical and atypical motion perception. In particular, integrating cortico-cortical neural noise may help clarify how noise contributes to motion detection deficits observed in schizophrenia. By bridging these gaps, this work offers a compelling framework for advancing the understanding of motion detection and its related impairments.

### 5.1 Decomposing the Motion Structure into Shared and Individual Motion Sources

Testing the model with various stimuli highlights the significance of incorporating a surround suppression mechanism into the motion detection process. Classical predictive coding provides some degree of surround suppression, but it fails to fully capture key characteristics of extra-classical receptive field effects necessary for accurately interpreting the complex interplay between object motion and its neighbors. Neurophysiological studies show integrative and antagonistic effects, where motion in the surround influences neuronal activity (Braddick, 1993; Zarei Eskikand et al., 2020; Eskikand et al., 2023). In the antagonistic case, neurons increase their activity when motion in the center and the surrounding area occur in opposite directions. Conversely, in the integrative case, neurons suppress their activity under similar conditions (Braddick, 1993; Tadin et al., 2003, 2019; Eskikand et al., 2023; Born, 2000; Huang et al., 2007). Our method incorporates this integrative effect to offer a more nuanced and comprehensive model for analyzing motion interactions within dynamic scenes. Including this mechanism is crucial for decomposing the motion structure and extracting shared motion within the visual field.

The surround suppression mechanism evaluates motion in both the center and the surround of the receptive field to check for consistency in the visual field. It suppresses neuronal activity when motion in the center and the surround occur in different directions (Born, 2000; Huang et al., 2007; Zarei Eskikand et al., 2020). Studies demonstrate that the center-surround mechanism plays a pivotal role in figure-ground segregation, edge detection, and distinguishing the movement of two overlapping or nearby objects within the receptive field (Braddick, 1993; Tadin et al., 2003, 2019; Eskikand et al., 2023). This model underscores the importance of such mechanisms for detecting global (shared) motion in the visual field. Implementing the center-surround mechanism in the model, based on the predictive coding algorithm, demonstrates this critical function effectively.

We tested the model using three distinct types of stimuli known for their significant contributions to the study of global motion detection in the visual cortex: (I) the Johansson experiment, (II) Duncker wheel, and (III) random dot kinematogram. The Johansson experiment I shows how the brain integrates information from multiple points into a coherent perception of global motion. The Duncker wheel experiment II demonstrates the tendency of our brain to break down multiple complex movements into recognizable, coherent motions, such as that of a rotating wheel in this experiment. The random dot kinematogram experiment III allows motion coherence to be manipulated by changing the level of noise to study how the brain integrates or segregates motion signals from different spatial locations. The proposed model effectively decomposes motions within the visual field into shared and individual object motions across all these stimuli, replicating the human brain’s perception of global motion (see Figure. 6b and Figure. 6d). By implementing a predictive coding algorithm, the model illustrates how the brain predicts an object’s future position by using its current and past states of motion, aligning with kinematic principles.

### 5.2 Balancing sensory data and prior knowledge in the detection of coherent motion and the relationship to schizophrenia

The modeling results demonstrate that increasing noise levels in the random dot kinematogram experiment III delays the model’s convergence time for detecting the global motions of the dots (see Figure. 8a). This finding replicates the psychophysics experiment bySpencer et al. (2013), which showed that participants need a longer duration of stimuli to detect global motion at higher noise levels. Their experiment compared healthy controls with patients with schizophrenia in detecting coherent motion in random dots. They found that patients with schizophrenia required longer durations than healthy participants to detect global motion across varying noise levels.

Using our model, we simulated the observed deficiency in coherent motion detection reported in patients with schizophrenia. The model indicates that increasing the influence of prior knowledge during motion detection significantly affects the convergence time. Extending the duration of the window for using past information in motion detection further prolonged the model’s convergence time, as shown in Figure. 8a. This simulation aligns with the observed differences in stimulus duration required by patients with schizophrenia compared to healthy controls for detecting coherent motion.

We analyzed this phenomenon further by varying the window’s duration for utilizing past information from 0 to 100 ms. The results revealed that increasing the window duration to a minimum threshold of 30 ms optimizes the model’s performance and reduces the time to convergence. Extending the window duration beyond this 30 ms threshold gradually lengthens the convergence time and ultimately causes the model to fail to converge, as illustrated in Figure. 8b. This finding highlights the importance of balancing prior knowledge and incoming sensory data in detecting coherent motion. The results demonstrate that the model must avoid overly relying on past information (i.e., a very large window duration) to detect coherent motion effectively. For optimal performance, the model requires some past information to prevent it from responding excessively to noise. Reducing the influence of past information (i.e., a very small window duration) increases the model’s convergence time, as the noise in the stimuli overly disturbs its detection of coherent motion.

These results suggest a possible explanation for the motion detection deficiency observed in patients with schizophrenia. Studies show that distortions in perception and beliefs among individuals with schizophrenia arise from accurately encoding prior knowledge, which makes them less responsive to feedback (i.e., under-learning) (Corlett et al., 2019; Horga and Abi-Dargham, 2019; Nassar et al., 2021). Other theories propose that abnormalities in motion perception result from interpreting prediction errors as significant changes, leading to exaggerated responses to noisy information and rapid adaptation to a changing environment compared to controls (i.e., over-learning) (Kaplan et al., 2016; Stephan et al., 2016). A recent study reveals the coexistence of these conflicting theories, showing that the learning process in these patients is not stationary and depends on the statistical context. The applied task demonstrated both over-learning and under-learning in different situations (Nassar et al., 2021). The model developed here indicates that the motion detection deficiency in patients with schizophrenia arises from the same mechanism, either by overly relying on past information (under-learning) or by excessively updating based on feedback (over-learning).

### 5.3 Surround suppression and the detection of coherent motion

The results of this study demonstrate how surround suppression affects the detection of coherent motion. Increasing the threshold for surround suppression improves the model’s performance in detecting coherent motion. Conversely, decreasing the threshold for surround suppression suppresses neuronal activity and increases the model’s convergence time. These findings suggest designing an experiment to test the surround suppression mechanism in patients with schizophrenia compared to controls. Such an experiment could offer new insights into the neural circuitry underlying surround suppression and how it is altered in patients with schizophrenia.

### 5.4 Comparisons with existing computational models of motion perception

The model developed by Bill et al. (2022) and Gershman et al. (2016) offers valuable insights into the hierarchical inference of visual motion structure by using a Bayesian framework to represent the relationships between multiple motion components. Their approach integrates sensory inputs with high-level probabilistic inference to explain how the brain extracts coherent motion from complex visual stimuli. Our model, grounded in the predictive coding framework, pursues the same objective of representing hierarchical motion perception. However, our model explicitly incorporates a surround suppression mechanism to simulate the neurobiological process of motion integration, which modulates motion signals based on their contextual relationship with surrounding stimuli. This mechanism plays a critical role in distinguishing between shared and individual motions in dynamic environments, an aspect not explicitly addressed in the model by Bill et al. (2022) and Gershman et al. (2016). Furthermore, while their model captures motion structure through Bayesian inference, our model also leverages the balance between prior knowledge and incoming sensory data to replicate motion detection processes, particularly those related to motion perception deficiencies in schizophrenia. By focusing on pathological aspects of motion detection, our approach introduces a novel perspective that differentiates it from other models regarding its clinical research applications.

Classical predictive coding models, such as that proposed by Rao and Ballard (1999), explain several visual cortical responses, including extra-classical receptive-field effects. However, they do not fully address motion perception, which introduces additional complexities. Although Rao and Ballard’s model includes feedback for spatial prediction, it does not account for the dynamic aspects of motion stimuli or the role of surround suppression in motion detection. Our model addresses these limitations by extracting motion structure and accounting for interactions between center and surround motion signals, which are critical for accurate motion detection. Our model demonstrates that motion perception under noisy conditions requires mechanisms that integrate motion consistency and surround suppression to filter irrelevant motion signals. Unlike classical models, our approach emphasizes the temporal dynamics of motion and the importance of context-dependent modulation, particularly in scenarios where global motion is disrupted.

Recent models like the dynamic predictive coding (DPC) framework introduced by Jiang and Rao (2022), enhance our understanding of spatiotemporal prediction in the neocortex. Their model organizes sequence learning hierarchically, where lower cortical levels predict short-term dynamics and higher levels generate abstract representations across longer timescales. This multi-level approach captures predictive and postdictive effects, offering insights into phenomena like the flash-lag effect (Nijhawan, 1994). While the DPC framework explains general hierarchical sequence processing and episodic memory formation, our model integrates surround suppression and predictive coding to simulate motion perception impairments under pathological conditions, such as schizophrenia. Our model emphasizes how alterations in motion signal integration, particularly through surround suppression, contribute to deficits observed in patients with schizophrenia. By focusing on clinical phenomena like increased latency in motion detection and over-reliance on past information, our model complements the DPC framework while providing a specialized focus on perceptual abnormalities in mental disorders.

Various computational models address schizophrenia by modeling neural circuitry, synaptic dysfunctions, and perceptual anomalies (Valton et al., 2017). However, most models overlook specific motion detection deficiencies in patients with schizophrenia. Existing models typically concentrate on higher-order cognitive deficits, reward-based learning, and hallucination-like symptoms, with few addressing sensory-level impairments like motion perception. Our model fills this gap by simulating impaired motion perception in schizophrenia patients due to aberrant neural dynamics. This focus on low-level sensory processing anomalies represents an unexplored domain in computational psychiatry, complementing existing models of higher-order cognitive dysfunctions in schizophrenia research.

To our knowledge, no existing model specifically addresses motion detection deficiencies in schizophrenia within the predictive coding framework. Previous studies link predictive coding impairments to broader aspects of schizophrenia, such as cognitive and perceptual disruptions (Liddle and Liddle, 2022). However, our study is the first to focus explicitly on how these impairments manifest in motion detection.

## Acknowledgment

This research is funded by Early Career Research Grant (ECRG) from the University of Melbourne.

## Appendix

We investigated the effect of the number of dots on the model’s computation and convergence time, Figure. 11. The results show that while the number of dots impacts convergence time, the effect diminishes as dot density increases. Convergence time decreases and eventually stabilizes. This supports our hypothesis that the model’s performance is robust due to its reliance on predictive coding and surround suppression mechanisms rather than the absolute number of dots.

**Figure 11:**
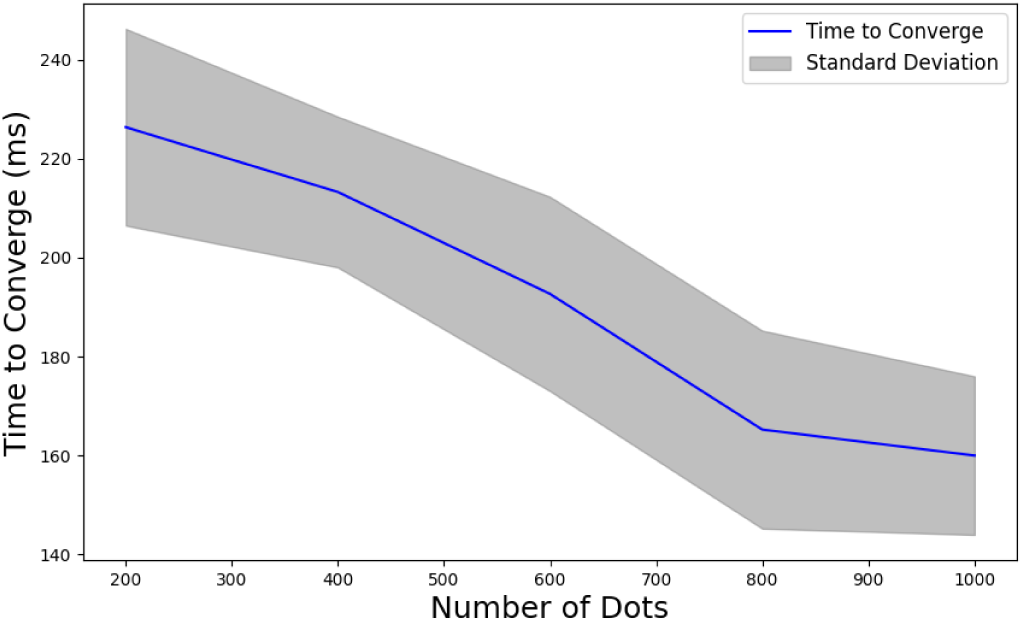
Effect of Dot Density on Convergence Time in Coherent Motion Detection. The blue line represents the mean convergence time, while the shaded gray region indicates the standard deviation.

## Notes

### Competing Interest Statement

The authors have declared no competing interest.

### Summary of Updates

This version of the manuscript has been revised to update the introduction, description of the methods and also discussion.

